# Effects of Covering Mature Avocado ‘Pinkerton’ Trees with High-Density Shading Nets during Cold Winters on Microclimate, Chlorophyll Fluorescence, Flowering and Yield

**DOI:** 10.1101/2023.07.03.547459

**Authors:** Lior Rubinovich, Carmit Sofer-Arad, Simon Chernoivanov, Nitzan Szenes

## Abstract

Avocado (*Persea americana* Mill.) is a subtropical fruit tree of high commercial value with increasing global demand. Most avocado cultivars are vulnerable to cold climates, which may reduce yields and restrict their geographical expansion. This includes the green-skinned avocado cv. Pinkerton, which accounts for 45% of the avocado cultivated in northeastern Israel. Shading nets can protect agricultural crops from cold environments. We therefore evaluated the effect of covering mature ‘Pinkerton’ trees with high-density shading nets during the winter. Trees were covered with Silver-coloured 50% or 70% shading nets during three consecutive winters, while uncovered trees served as controls. Photosynthetically active radiation in plots covered with the Silver 50% or 70% nets was significantly lower than for the control by 52% and 90%, respectively. Minimum air temperature was similar between treatments. Maximum air temperature was generally lower under the shading nets compared to the control. The ratio of variable to maximum fluorescence (Fv/Fm) measured in February 2019 and 2020 was 0.72 and 0.8 in the control trees, 0.79 and 0.83 in the Silver 50% trees and 0.81 and 0.84 in the Silver 70% trees, respectively. Flowering intensity was lower in the net-covered trees compared to the control, by up to 42%. Interestingly, the three-year average yield of trees covered with the Silver 50% or 70% nets was insignificantly higher by 27% and 38%, respectively, compared to the control trees. These results suggest that the reduction of daytime solar irradiance in the winter by the shading nets may mitigate cold stress and increase yield. Additional long-term studies should examine the effects of shading nets and other shading strategies on different avocado cultivars.

## Introduction

Avocado (*Persea americana* Mill.) is a subtropical fruit tree of high commercial value with increasing global demand. In 2021, avocado world production was ∼8.7 million tons with a gross production value of ∼8.56B $US (www.fao.org, accessed 7.2023). Most avocado cultivars are vulnerable to cold stress, which may reduce yields and restrict their geographical expansion (Schaffer et al. 2013). Extreme cold events are becoming more frequent as climate change intensifies, even in regions where they were previously rare. As a result, it is imperative to develop solutions to reduce cold damage in mature avocado plantations. The green-skinned avocado cv. Pinkerton is considered as cold-sensitive as ‘Hass’ and accounts for 45% of the avocado cultivated in northeastern Israel. The majority (∼70%) of the Israeli ‘Pinkerton’ are exported to Europe (mainly Eastern Europe), while ∼30% are marketed in the domestic market (data from the Israeli Fruit Board). In this region, minimum winter temperatures may drop below 0 °C (frosts) or approach, but remain higher than 0 °C (chilling events). Whereas frost can cause mild to severe visible damage to leaves, buds, flowers and fruit, the consequences of chilling have been much less described, because the plant tissues show almost no visible damage (Chernoivanov et al. 2022). Chilling temperatures can cause photoinhibitory damage to photosystem 2 (PS II) that can be quantified by measuring a decrease in the ratio of variable to maximum chlorophyll fluorescence (Fv/Fm). Whiley et al. 1999 showed that in avocado and mango low winter temperatures below 10° C combined with high light intensity resulted in a decrease in Fv/Fm values due to chill-induced photoinhibition of leaves (Whiley et al. 1999). Additional studies showed that Fv/Fm rate is a reliable quantitative indicator of cold stress damage (Rizza et al. 2001). Shading nets can protect agricultural crops from harsh environments, including cold damage (Chernoivanov et al. 2022; Zait et al. 2020). We hypothesized that covering avocado trees during the winter with high-density shading nets would positively affect the trees and reduce cold damage. Therefore, the main objective was to evaluate the effect of covering mature ‘Pinkerton’ trees with different shading nets during the winter. Specifically, we evaluated the capacity of the shading nets to reduce the risks of cold damage and to increase tree production.

## Materials and Methods

### Experimental site

The experiments were conducted during 2017-18, 2018-19 and 2019-20 in a commercial ‘Pinkerton’ orchard in Kibbutz Dan in northwest Israel (lat. 33°23’N, long. 35°66’E, 190 m above sea level). ‘Ettinger’ trees served as pollinizers. The trees were grafted on ‘Shiller’ seedling rootstocks and planted in 1996 with 6-m spacing between rows and 4 m between trees. The rows were oriented northeast/southwest. Chilling temperatures occurred in the experimental orchard mainly from December to mid-February.

### Shading nets and experimental design

From early December through mid-March (before the flowering period) of 2018, 2019 and 2020, Silver-coloured nets with 50% or 70% shading level (Silver 50% and 70%, Ginegar Plastic Products Ltd., Ginegar, Israel) were placed over the trees, supported by their canopies. Trees from the control plot were not covered with shading nets (control trees). The experimental plots were completely randomized, with four repeats for each treatment (control and shading nets, n=4). The area of each repeat was 0.1 ha, and measurements were taken only from the eight middle trees of each repeat (experimental trees).

### Temperature and light measurements

Miniature waterproof, single-channel Hobo temperature data loggers (cat. no. UA-001-64; Onset Corp., Bourne, MA, USA) were used to measure air temperatures in the field. They were positioned 1.5 m above the ground and shielded from direct sunlight by the canopy. The air temperature was measured continuously at 10-min intervals from December until mid-March of each year. Three temperature data loggers were placed for each treatment. Photosynthetically active radiation (PAR) was measured 1.5 m above the ground (∼3 m beneath the nets) using a Field Scout quantum meter (#3415F; Spectrum Technologies Inc., Aurora, IL, USA). At least 10 measurements were performed for each treatment with the different nets and the controls.

### Analysis of chlorophyll a fluorescence

Chlorophyll a fluorescence was measured shortly before sunrise in complete darkness with a FluorPen FP100 portable fluorometer (Photon Systems Instruments, Drasov, Czech Republic) and Fv/Fm was calculated (Chernoivanov et al. 2022). Measurements were taken during winter in Feb 2019 and 2020, following the swelling of inflorescence buds and before flowering, from at least 3 leaves per each of the experimental trees.

### Estimation of flowering intensity and yield measurements

Flowering intensity was appreciated as described previously during peak bloom in Apr 2018, 2019 and 2020 (Ziv et al. 2014); in a blind test, two surveyors independently assessed each experimental tree from 0 (no flowering) to 5 (high flowering intensity). To determine fruit yield, during the commercial harvest period in each year (winter), all the fruits on the experimental trees were manually picked and weighed separately for each tree. The three-year mean yield of each treatment was calculated by averaging the yields of each of the three years of the experiment (n = 3).

### Statistical analysis

All results were subjected to one-way ANOVA followed by Tukey-HSD test using JMP software, version 11.0.0 (SAS Institute, Cary, NC, USA).

## Results

### Effect of shading nets on PAR and air temperature

PAR in plots covered with the Silver 50% or 70% nets was significantly lower (*P* <0.001), by 52% and 90% compared to the control, respectively (Supplementary Fig. 1). Thus, the actual shading rate of the Silver 70% shading nets was slightly higher than expected. PAR in plots covered with the Silver 50% net was significantly higher (*P* <0.001) than in those covered with the Silver 70% net. During the winters of 2017-18 and 2018-19, the minimum air temperature in the orchard did not fall below 0°C, reaching 2.52°C (Fig. 1A) and 0.42°C (Fig. 1B), respectively. In the winter of 2019-20, minimum air temperature reached below 0°C (−1.38°C), but on only one night (Fig. 1C). Minimum air temperature was similar between the treatments (Fig. 1A,B,C). However, maximum air temperatures under the shading nets was generally lower compared to the control (Fig. 1D,E,F)

**Fig. 1.**
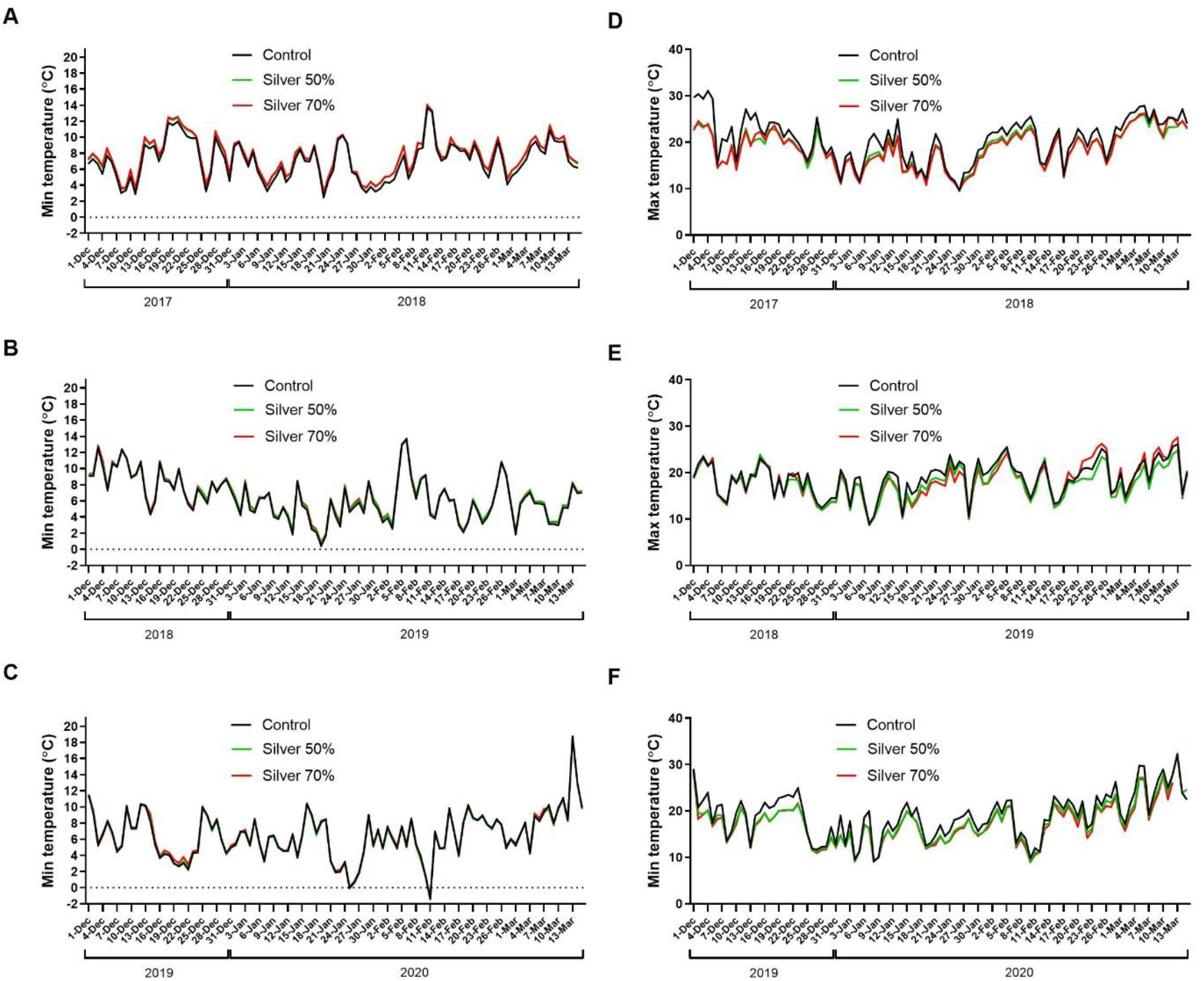
Minimum and maximum daily air temperatures in the control plots and under the different shading nets during the winters of 2017-18 (A,D), 2018-19 (B,E) and 2019-20 (C,F). The average of three data loggers represents the mean temperature for each treatment.

### Effect of shading nets on Fv/Fm

In Feb 2019 and 2020, Fv/Fm rate in the leaves of trees from the control treatment reached 0.72 and 0.8, respectively (Supplementary Fig. 2A). During both years, Fv/Fm rates were significantly higher (*P* < 0.05) under the Silver 50% and 70% shading nets, with no significant differences between them (*P* > 0.05).

### Effect of shading nets on flowering intensity and yield

The baseline (background) yield before the deployment of the nets was meager and similar between treatments (Supplementary Fig. 2B). In Apr 2018, flowering intensity was very high, with no significant differences (*P* > 0.05) between the treatments (Fig. 2A). In Apr 2019, flowering intensity in the control trees was high, significantly higher (*P* < 0.05) than that of the trees covered with the Silver 70% net. In Apr 2020, flowering intensity was highest in the control trees, but with no significant differences (*P* > 0.05) between the treatments. At the end of 2018, fruit yield was significantly higher (*P* < 0.05), by 71%, in the trees covered with the Silver 70% net compared to the control trees (Fig. 2B). There were no significant differences (*P* > 0.05) between the yields of trees covered with the Silver 50% net compared to both control and Silver 70%-covered trees. At the end of 2019, fruit yield was very low, with no significant differences (*P* > 0.05) between the treatments. However, the yields of trees covered with the Silver 50% or 70% nets were higher by 109% and 62%, respectively, compared to the control trees. At the end of 2020, average fruit yield was similar, with no significant differences between the treatments (*P* > 0.05). The three-year average yield of trees covered with the Silver 50% or 70% nets was insignificantly (*P* > 0.05) higher by 27% and 38%, respectively, compared to the control trees.

**Fig. 2.**
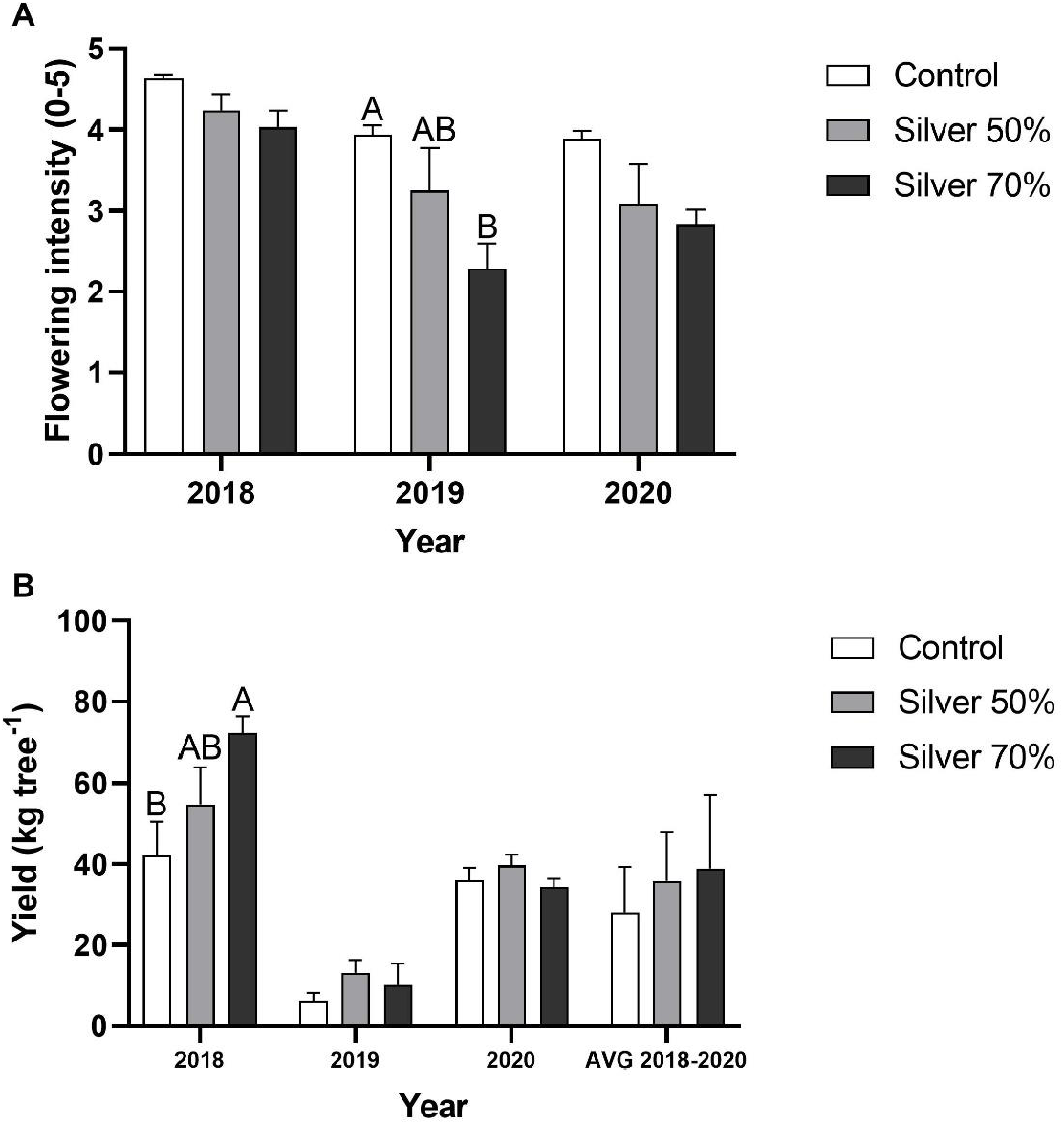
Flowering intensity was assessed and scored in Apr 2018, Apr 2019 and Apr 2020 on a scale of 0–5, with 0 representing no apparent flowering and 5 representing maximum bloom (A). During the commercial harvest period of each year, fruits were picked and weighed separately for each tree (B). Values are means ± SE of four repeats (n=4), each comprised of eight trees. Values of the three-year average yield (AVG 2018-2020) are means ± SE of the yields of each year of the experiment (n=3). Different letters above a column indicate significant differences within each year (Tukey-HSD, *P* < 0.05).

## Discussion

A decrease in Fv/Fm may indicate chill-induced photoinhibition and cold stress damage (Whiley et al. 1999; Rizza et al. 2001). Thus, the higher Fv/Fm values observed in trees covered with either of the shading nets compared to the control suggest a reduction in cold stress. A previous study showed similar results in young avocado ‘Reed’ trees covered with the same shading nets during cold winters, where temperatures were close, but above 0° C (Chernoivanov et al. 2022).

Flowering intensity was generally lower in the net-covered trees. Floral induction in avocado grown in the northern hemisphere occurs during late autumn, before net deployment (Ziv et al. 2014). Thus, it is possible that the reduction in flowering intensity in the net-covered trees was due to the reduction in winter daytime temperatures and/or solar irradiation, affecting floral bud development, which occurs during winter (Acosta-Rangel et al. 2021). This assumption requires substantiation. It is also possible that alternate bearing, which is manifested by a decrease in flowering intensity following high yield (Lovatt 2010), was the reason for the reduction in flowering intensity in the net-covered trees in April 2019. Still, during the years of the experiment, despite the lower flowering intensity there was no reduction in fruit yield with both net treatments. In particular, at the end of 2018, yield in the net-covered trees was higher than the control and was inversely related to flowering intensity of April 2018. Thus, it is possible that the use of these shading nets during winter enhances avocado flowering quality (i.e., the functionality of floral organs) rather than flowering intensity, leading, in some cases, to higher yields. It is also possible that the use of shading nets resulted in a delay in flowering, which may be advantageous to fruit yield in cases of low spring temperatures, where bee pollination is impaired (Stern et al. 2021). Covering the avocado trees during winter may also enhance chlorophyll content and photosynthesis rate, thus increasing carbohydrate production required for fruit set and development (Mditshwa et al. 2019, Alon et al. 2022, Chernoivanov et al. 2022). These possible effects were not examined in this study and therefore warrant further investigation.

The relatively low yields in 2019 following the high yields of the previous year can be attributed to the tendency of avocado to alternate bearing (Cohen et al. 2023). In light of the fact that the yield of the net-covered trees in 2018 was higher compared to the control, it could be expected that the yield in the following year in these trees would be lower compared to the control. The fact that the yield in the net-covered trees was higher, further supports the effect of the shading nets on fruit yield.

It is important to note that similar to previously reported results, the shading nets did not elevate minimum winter temperatures, but reduced daytime maximum temperatures and solar irradiation (Chernoivanov et al. 2022). High light intensity plays an important role in overall stress during chilling events (Wise 1995). Thus, the effect of the nets on Fv/Fm values and fruit yield might be attributed to the reduction of daytime solar irradiance in the winter, rather than changes in minimum air temperature. This assumption requires further investigation of the mechanism in which the shading nets exert their effects, for example by determining light-adapted chlorophyll fluorescence, leaf temperature and gas-exchange parameters.

On a broader scope, previous studies have shown that high-density shading nets can protect banana plants from frost damage (Zait et al. 2020). Interestingly, high-density shading nets also improve photosynthetic performance in mature ‘Pinkerton’ trees during extreme heat events (Alon et al. 2022). Thus, using shading nets in avocado orchards should be considered a potential measure to reduce a wide range of climate-related abiotic stresses. However, excess shading may have adverse effects. For instance, photosynthetic performance may decrease on days with regular fall temperatures (Alon et al. 2022). Furthermore, bee activity and pollination may be impaired in trees grown under shading nets (Mditshwa et al. 2019). Hence, long-term follow-up studies should be conducted to establish proper shading-management protocols using shading nets and other shading strategies, such as over-canopy solar panels, with different avocado cultivars and in countries with different light regimes.

## Conclusion

In conclusion, covering ‘Pinkerton’ trees with high-density shading nets during cold winters potentially have a negative effect on flowering, but a positive effect on chlorophyll fluorescence and fruit yield. Although the mechanism for these effects remains unclear, it is possible that the reduction of daytime solar irradiance is involved. Additional studies should be conducted to further examine their mechanism of effect and to evaluate their ability to reduce tree damage during extreme cold events, such as frosts.

## Acknowledgements

The authors thank the Avocado-GAL Corporation and the Israeli Fruit Board for financial support, and the Kibbutz Dan avocado team and Michael Noy for their invested effort in this study.

## Notes

### Competing Interest Statement

The authors have declared no competing interest.

## References Cited

Acosta-Rangel A, Li R, Mauk P, Santiago L, Lovatt CJ. 2021. Effects of temperature, soil moisture and light intensity on the temporal pattern of floral gene expression and flowering of avocado buds (Persea americana cv. Hass). Sci Hortic. 280:109940. https://doi.org/10.1016/j.scienta.2021.109940.

Alon E, Shapira O, Azoulay-Shemer T, Rubinovich L. 2022. Shading nets reduce canopy temperature and improve photosynthetic performance in ‘Pinkerton’ avocado trees during extreme heat events. Agronomy. 12(6):1360. https://doi.org/10.3390/agronomy12061360.

Chernoivanov S, Neuberger I, Levy S, Szenes N, Rubinovich L. 2022. Covering young ‘Reed’ avocado trees with shading nets during winter alleviates cold stress and promotes vegetative growth. Eur J Hortic Sci. 87(1):1–10. https://doi.org/10.17660/eJHS.2022/007.

Cohen H, Bar-Noy Y, Irihimovitch V, Rubinovich L. 2022. Effects of seedling and clonal West Indian rootstocks irrigated with recycled water on ‘Hass’ avocado yield, fruit weight and alternate bearing. N Z J Crop Hortic Sci. :1–13. https://doi.org/10.1080/01140671.2022.2098779.

Lovatt CJ. 2010. Alternate Bearing Of ‘Hass’ Avocado. California Avocado Society Yearbook. 93:125–140.

Mditshwa A, Magwaza LS, Tesfay SZ. 2019. Shade netting on subtropical fruit: Effect on environmental conditions, tree physiology and fruit quality. Sci Hortic. 256:108556. https://doi.org/10.1016/j.scienta.2019.108556.

Rizza F, Pagani D, Stanca AM, Cattivelli L. 2001. Use of chlorophyll fluorescence to evaluate the cold acclimation and freezing tolerance of winter and spring oats. Plant Breeding. 120(5):389–396. https://doi.org/10.1046/j.1439-0523.2001.00635.x.

Schaffer B, Gil P, Mickelbart M, Whiley A. 2013. The avocado: Botany, production and uses. CABI, Wallingford. https://doi.org/10.1079/9781845937010.0000.

Stern RA, Rozen A, Eshed R, Zviran T, Sisai I, Sherman A, Irihimovitch V, Sapir G. 2021. Bumblebees (Bombus terrestris) improve ‘Hass’ avocado (Persea americana) pollination. Plants. 10(7):1372. https://doi.org/10.3390/plants10071372.

Whiley AW, Searle C, Schaffer B, Wolstenholme BN. 1999. Cool orchard temperatures or growing trees in containers can inhibit leaf gas exchange of avocado and mango. J. Am. Soc. Hortic. Sci. 124(1):46–51. https://doi.org/10.21273/jashs.124.1.46.

Wise RR. 1995. Chilling-enhanced photooxidation: The production, action and study of reactive oxygen species produced during chilling in the light. Photosynth Res. 45(2):79–97. https://doi.org/10.1007/BF00032579.

Zait Y, Elingold I, Londener A, Gal E, Or G, Galpaz N. 2020. Banana frost protection by thermal nets, p 21–26. Acta Horticulturae. International Society for Horticultural Science. https://doi.org/10.17660/ActaHortic.2020.1272.3.

Ziv D, Zviran T, Zezak O, Samach A, Irihimovitch V. 2014. Expression profiling of FLOWERING LOCUS T-like gene in alternate bearing ‘Hass’ avocado trees suggests a role for PaFT in avocado flower induction. PLoS One. 9(10):1–14. https://doi.org/10.1371/journal.pone.0110613.

