## Supplemental material for "Effects of Covering Mature Avocado ‘Pinkerton’ Trees with High-Density Shading Nets during Cold Winters on Microclimate, Chlorophyll Fluorescence, Flowering and Yield"

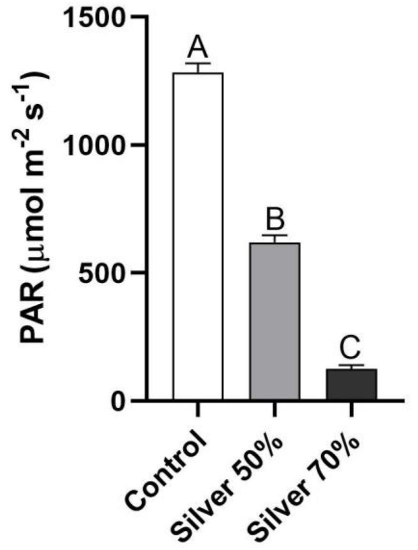


Supplementary Fig. 1. Effect of winter shading nets on photosynthetically active radiation (PAR). PAR measurements were taken in Feb 2018 during the late morning in the control and net-covered plots. Values are means ± SE of 10 replicates ($n = 10$). Different letters above columns indicate significant difference (Tukey-HSD, $P < 0.05$).

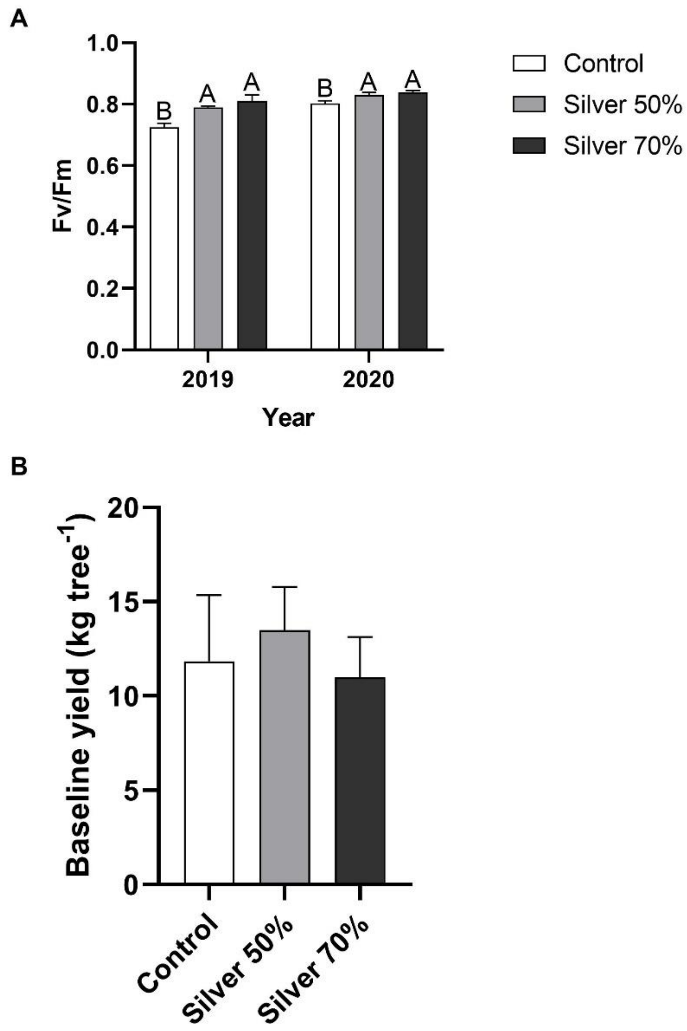


Supplementary Fig. 2. Chlorophyll a fluorescence parameters were recorded in Feb 2019 and Feb 2020 in the dark shortly before sunrise to calculate maximum quantum yield (Fv/Fm, A). Values are means ± SE of four repeats (n=4), each comprised of eight trees, three leaves were measured from each tree. On January 2018 (the commercial harvest of season 2017-18, baseline), fruits were picked and weighed (B). Values are means ± SE of four repeats (n=4), each comprised of eight trees. Different letters above columns indicate significant difference (Tukey-HSD, $P < 0.05$).
